# CAPRINI-M: An AI-curated Cardiac-Specific Atlas of Protein Interactions in Mice

**DOI:** 10.64898/2026.03.06.710104

**Authors:** Enio Gjerga, Philipp Wiesenbach, Chris-Andris Görner, Ying Zhang, Konstantin Pelz, Markus List, Christoph Dieterich

## Abstract

**Motivation:** Protein–protein interactions are fundamental to cardiovascular disease biology, but the corresponding knowledge is dispersed across the literature and heterogeneous databases, making systematic curation time-consuming. Moreover, many existing PPI resources may be biased and lack detailed information on structural interaction interfaces or associated thermodynamic parameters.

**Results:** We present CAPRINI-M (CArdiac PRotein INteractions In Mice), a web-based tool hosting an AI-curated atlas of cardiac protein interactions. We mined 9,105 cardiobiology manuscripts and used open-source LLMs (LLaMA-3.3 70B) to extract 11,189 protein–protein interactions. We then used AlphaFold3 to infer interaction interfaces, estimate thermodynamic properties related to complex stability, and predict the likelihood that each protein pair forms a complex. In our benchmarking analysis, CAPRINI-M showed stronger performance than the comparator PPI resources tested here. Predicted interaction favourability also agreed with published experimental evidence, with lower predicted Gibbs free energy associated with experimentally preferred binding partners. Overall, CAPRINI-M provides a more comprehensive, mechanistically annotated view of cardiovascular disease–relevant protein–protein interactions by integrating literature evidence with structural, interface-level, and stability-related information.

**Availability:** The CAPRINI-M web application is available at https://shiny.dieterichlab.org/app/caprinim. The source code used in this study is linked in the manuscript’s Availability section.

## 1. Introduction

Protein–protein interactions (PPIs) play vital roles in the emergence and progression of cardiovascular diseases (CVD) (Wheeler-Jones, 2005). Disruptions in normal PPI networks may contribute to the development of key processes in CVDs, including cardiac remodelling, inflammation, dysfunctional calcium handling, and heart failure (He et al. 2022).

PPI databases provide a useful foundation for functional and mechanistic studies of cardiac signalling pathways. Many general-purpose PPI databases have been established over the years, providing a valuable backbone for studying signalling pathway systems (Türei et al. 2021; Szklarczyk et al. 2022; Oughtred et al. 2021; del Toro et al. 2021; Xenarios et al. 2000). However, these resources typically rely on manual curation, which is labour- and time-intensive, and maintaining them in an up-to-date state is challenging given the continuously expanding literature. In this regard, the emergence of Large Language Models (LLMs) has unlocked new opportunities to mine domain-specific molecular interactions from the literature. These models, while not necessarily trained exclusively on scientific literature, can readily extract knowledge in the form of entity relations across scientific domains (Zhao et al. 2024). At the same time, large-scale LLM-based relation extraction may not be reliable as prompting can introduce hallucinations (relations between absent entities), false negatives (missed interactions), and false positives (incorrect interactions between present entities). This makes it critical to quantify extraction performance using curated benchmarks before deploying LLM-driven mining as a backbone for biological interpretation. Another issue is that general PPI resources tend to be biased towards over-represented domains, such as cancer (Blumenthal et al. 2024). This may complicate their use for studying cardiac systems, as it may introduce bias toward proteins and pathways that reflect better-characterised domains rather than cardiac-specific biology. Moreover, not all resources, with DIGGER as a notable exception (Albrecht et al. 2025), provide information about the protein interfaces involved in the interaction. This is an especially important aspect to consider when studying signalling systems involved in CVDs, as alternative splicing (AS) – a critical post-transcriptional regulator of many cardiac mechanisms (Beqqali 2018) – can remodel PPI networks by altering, adding, or removing interaction-relevant regions. Building on this, our previous work introduced LINDA (Linear Integer programming for Network reconstruction using transcriptomics and Differential splicing data Analysis) (Gjerga et al. 2023), a mechanistic framework that integrates transcriptomics, differential splicing evidence, and domain-centric interaction knowledge to infer splicing-dependent rewiring of regulatory and protein interaction networks. While LINDA can capture important cases where splicing removes or adds entire exons/domains, many biologically relevant events are subtler (e.g., alternative splice sites, microexon insertions, partial exon truncations, or alternative N-/C-termini) and may perturb only part of an interface rather than eliminate an entire domain. Given this, automated and efficient extraction of PPIs has the potential to significantly improve the speed and precision of creating domain-specific knowledge resources aligned with the latest cardiovascular biology literature. Additionally, the most recent computational methods for structural prediction of proteins, such as AlphaFold3 (Abramson et al. 2024) or RosettaFold (Baek et al. 2021), offer unique opportunities to study the structures adopted by proteins in their interacting bound state. By analysing the static structures of these PPIs, we aim to accomplish two complementary objectives: first, to delineate with precision the specific regions or domains that mediate the interaction; and second, to approximate the thermodynamic properties of the resulting complex, thereby gaining insight into its stability and degree of transience. An interface-resolved perspective is particularly important, as many disease-associated variants and therapeutic interventions act by perturbing discrete binding surfaces rather than abolishing entire proteins or domains. Emphasising this interface-centric view also expands the range of splicing-related effects that can be detected and interpreted. Instead of being limited to complete exon or domain loss, we can capture partial or localised sequence alterations that selectively remodel binding interfaces and modulate interaction strength. Nevertheless, it remains non-trivial to determine which structural/interface signals are most informative, and how confidently one can infer that an extracted protein pair truly forms a stable interaction. This motivates combining literature-derived candidates with downstream modelling and learning-based assessment of interaction plausibility rather than treating all extracted edges as equally valid.

In this study, we present CAPRINI-M (CArdiac PRotein INteractions In Mice, https://shiny.dieterichlab.org/app/caprinim), an end-to-end, AI-powered framework accompanied by a web resource that transforms scattered cardiobiology literature into a structurally and thermodynamically annotated, cardiac-specific interaction knowledge base (**Fig. 1**). CAPRINI-M implements a comprehensive linear workflow that: ***(i)*** gathers and filters relevant literature; ***(ii)*** extracts and ensures the quality of interaction evidence; ***(iii)*** constructs a cardiac PPI atlas; ***(iv)*** structurally models each candidate complex; ***(v)*** quantifies properties related to interfaces and stability; and ***(vi)*** integrates all evidence into a searchable database and interactive web application.

**Fig. 1.**
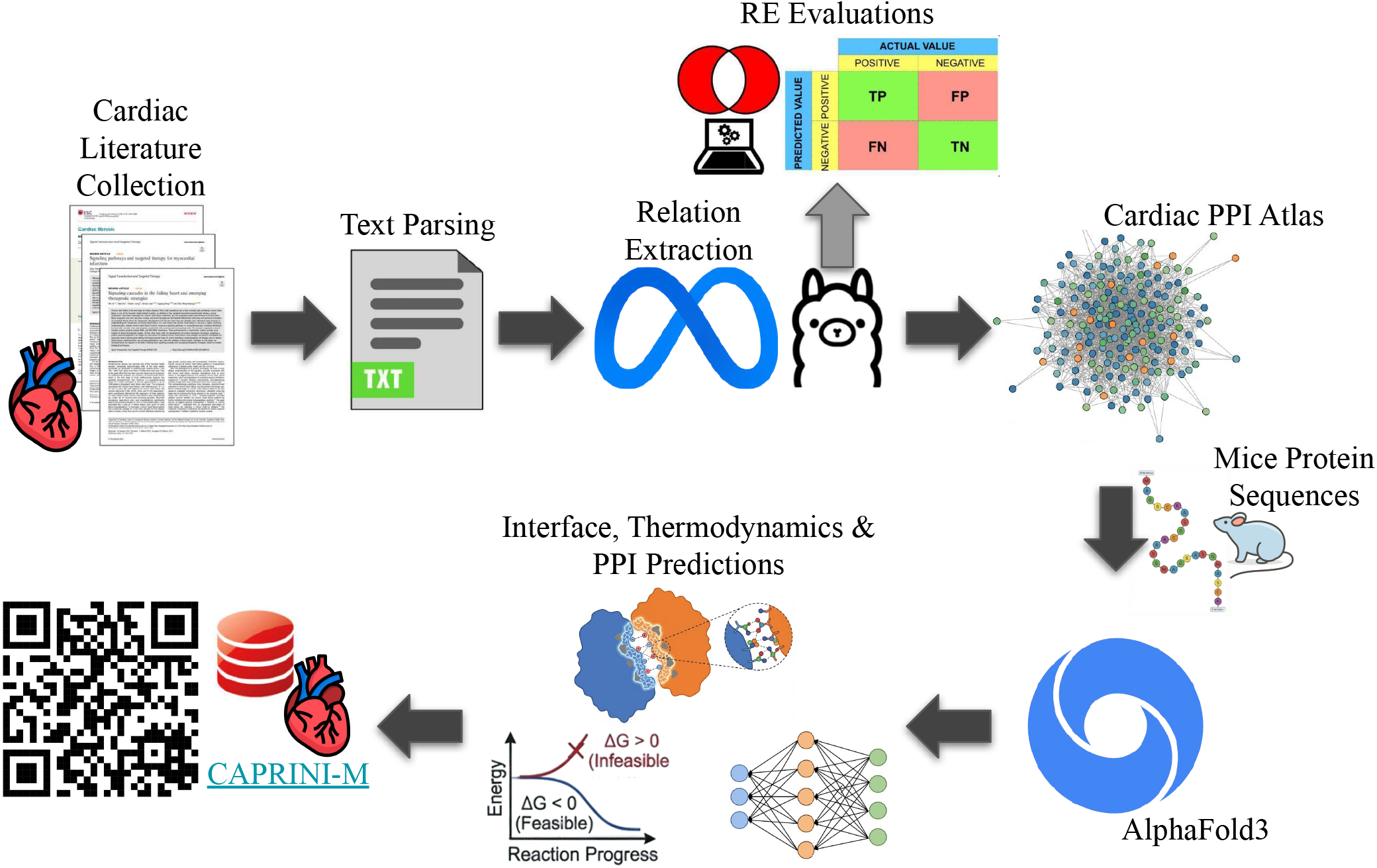
Cardiac manuscripts were fetched from PubMed Central and parsed into text files. Optimised prompt-based LLMs were used to extract PPIs from the text and generate a cardiac PPI atlas. Mice protein sequences were obtained for each protein present in the atlas, and PPI 3D structures were predicted using AlphaFold3. From 3D structures, we have identified interaction and thermodynamic features, as well as estimated interaction probabilities, using independently trained NN-based machine learning models. The mouse PPI atlas, along with all evaluations, is available in the CAPRINI-M web application.

As illustrated in the stepwise workflow in **Fig. 1**, CAPRINI-M begins by collecting cardiac-related literature from PubMed Central, after which the manuscripts are converted into text to enable scalable downstream analysis. Automated cardiovascular-domain screening is then applied to each manuscript, classifying papers as either cardiac-relevant or non-cardiac. This ensures that relation extraction is performed on a corpus specifically enriched for cardiac biology, avoiding dilution by broader biomedical topics. Next, relation extraction is carried out using open-source LLMs (LLaMA-3.3 70B) with optimised prompting strategies to recognise protein entities and extract candidate PPI statements. Additional normalisation steps are included to address name and synonym variability across different papers. To ensure reliability in large-scale LLM-based mining, we rigorously evaluate relation extraction performance (true/false positives and negatives) using a curated, annotated benchmark corpus before deploying the chosen strategy at scale. This approach allows us to quantify expected error patterns in the extraction stage.

The extracted relations are then unified into a cardiac PPI atlas for mouse, which serves as a literature-backed foundational network. From this atlas, we retrieve the corresponding mouse protein sequences for each interaction partner and submit every protein pair to AlphaFold3 to predict the structure of the bound complex. Using these predicted 3D complexes, we generate interface descriptors (detailing where and how proteins interact) and thermodynamic or stability indicators (such as ΔG estimates). This moves beyond simply identifying connections, providing deeper mechanistic insights that enable users to analyse binding surfaces, potential partner competition, and the impact of sequence changes on interaction strength. Finally, we evaluate the extracted PPIs by training and applying AI classifiers that combine AlphaFold3-derived interface and quality metrics with sequence-embedding features (ESM2/ESM3/ESMC (Lin et al. 2023; Hayes et al. 2025; ESM Team, 2024) to estimate the likelihood of interaction. This allows us to categorise pairs as more or less likely to form feasible complexes. All components (literature evidence, extraction diagnostics, atlas structure, predicted models, interface and thermodynamic annotations, and interaction-likelihood scores) are integrated into the CAPRINI-M database, which is accessible through an interactive web application for searching, filtering, and viewing structures in detail.

The CAPRINI-M resource, with its multiple features, is available online (https://shiny.dieterichlab.org/app/caprinim). A key hypothesis motivating CAPRINI-M is that a cardiac-focused, literature-derived PPI atlas should act as a more appropriate backbone network for cardiovascular systems analyses than general-purpose PPI resources, which may be biased toward better-studied non-cardiac domains. If this is true, then downstream analyses that depend on network structure, such as module reconstruction from disease omics data and topology-aware pathway enrichment, should more strongly and more specifically recover expected gene sets when using CAPRINI-M than when using a standard, general interactome. In this work, we demonstrate the value of CAPRINI-M in recapitulating concise cardio-focused network/module reconstruction and pathway edge-topology enrichment analyses, and in accurately predicting which experimentally verified competing interactions are more favourable.

## 2. Methods

### 2.1 AI-Based Relation Extraction (RE)

To identify the most suitable prompting strategy for large-scale PPI extraction from our cardiovascular manuscript corpus (Section 2.2), we first benchmarked multiple prompt-engineering variants on the annotated RegulaTome corpus (Nastou et al. 2024). RegulaTome contains 2,521 short scientific documents and 16,962 annotated relations. After mapping relation types into categories, the PPI subset of the RegulaTome comprised 6,435 PPIs (Train: 3,870; Validation: 1,301; Test: 1,264 – from the original defined RegulaTome data split) (Supplementary Text 1). We then implemented a prompt-based RE workflow in which the LLaMA-3.3 70B model (Siriwardhana et al. 2024) was instructed to output extracted relations and RegulaTome text ID in JSON format. We evaluated our strategy under two conditions: LLM-only (document text only) and LLM–spaCy (document text plus entity names extracted with spaCy and provided as contextual guidance; Supplementary Figure 1; Yenkikar et al. 2024). The prompts used for PPI RE have been summarised in Supplementary Text 2. We subsequently evaluated a set of complementary prompt variants designed to address common error types in document-level relation extraction. Example-based prompting (*PosExamples, NegExamples, AllExamples*) was employed to guide the LLaMA-3.3 70B model: positive examples guide the model on how to identify PPIs from the text, while negative examples are used to reduce frequent false positives (e.g., co-expression or association statements). The *AllExamples* strategy combines the base prompt with *PosExamples* and *NegExamples*. Retrieval-augmented in-context examples (*DynExamples*) adapt the prompt to each manuscript’s writing style by incorporating the most similar curated examples. External partner-context augmentation (*STRINGLookup*) provides additional biological context for candidate partners, minimising ambiguity in protein mentions and facilitating accurate pairing. Finally, robustness and consensus strategies (*ToT-N3, Ensembl-N5*) (Dyer et al. 2025), where we repeat extraction multiple times (either via three reasoning paths (ToT-N3) or five independently sampled outputs (Ensembl-N5)) and then aggregate/keep interactions that recur. Each strategy was assessed using the annotated RegulaTome corpus to identify the best-performing approach, and then applied to large-scale PPI extraction in the cardiovascular corpus. PPIs were treated as undirected edges to avoid directionality artefacts, and to mitigate entity string variation from LLM outputs, we applied synonym expansion (up to 10 synonyms) by using LLaMA-3.3 70B with a dedicated synonym prompt (Supplementary Text 2D). We then matched predictions to the ground truth by accepting a PPI as correct if either orientation of the pair matched and if any synonym (including the original name) for each interactor matched the corresponding annotated entity. RE performance over the annotated RegulaTome corpus was reported as precision, recall, and F1, with uncertainty quantified via nonparametric bootstrapping (100 resamples) to compute 95% confidence intervals and variability statistics.

### 2.2 Cardiovascular Corpus

The RegulaTome annotated corpus introduced above serves as a reference standard for evaluating relation extraction (RE) performance and identifying the most effective prompting strategy. This optimised approach was then applied to curate cardiac full-text manuscripts, which we systematically mined to extract protein–protein interactions at scale for CAPRINI-M. The cardiac manuscript corpus was retrieved from PubMed Central (https://pmc.ncbi.nlm.nih.gov/) (see Supplementary Text 3A for details). Using this strategy, we obtained 9,105 manuscripts. A majority-vote, three-prompt-based strategy (Supplementary Text 3B) was then applied to the downloaded corpus to filter out outside of the cardiovascular domain. We retained 7548 scientific papers post-filtering. Each of the remaining manuscript PDFs was then parsed into Markdown text, and the best prompting strategy identified in our RegulaTome evaluations was applied to extract PPI relations. We evaluated 7548 papers, and 4503 of them reported at least one PPI. In this way, we identified and reported 11,189 PPI relations.

### 2.3 AlphaFold3 Evaluations

We implemented a high-throughput AlphaFold3 pipeline to model predicted protein–protein complex structures for our set of LLM-based PPI relations (see scripts/alphafold3_analysis_workflow in https://github.com/enio23/caprinim_ppi_info/). This pipeline batch-processes the list of extracted PPI pairs by running AlphaFold3 in two SLURM array stages, where it first takes each PPIs input protein sequence pairs and runs the data/alignment pipeline to generate the aligned outputs; then it locates those alignment-result directories and runs GPU inference to produce the final AlphaFold3 predictions and confidence summaries. For each PPI, five alternative 3D structures were generated, and we selected the best based on the highest internal confidence/ranking scores provided by AlphaFold3. For each PPI 3D structure, we extracted confidence estimates directly from AlphaFold3 and calculated our own metrics that reflect the strength of the PPIs and the interfaces mediating them.

### 2.4 Thermodynamic Properties

We evaluate the AlphaFold3-predicted protein–protein complexes by taking each complex structure, separating it into chains representing the interacting protein partners, generating AMBER (Pearlman et al. 1995) topology/coordinate files with a standard protein force field, running a brief energy minimisation, and then computing an approximate binding free energy using a single-frame MM/GBSA (MMPBSA.py) calculation (Milller et al. 2012). From this analysis, we report Van der Waals, Surface Energy, and Gibbs Free Energy (ΔG) approximates for each complex, enabling a consistent, high-throughput assessment of interaction stability/likelihood across many predicted PPIs.

Predicted binding free energies (ΔG) can be used to prioritise competitive protein–protein interactions. For each interaction edge, the reported ΔG value can be interpreted as a proxy for binding strength, with more negative ΔG values corresponding to thermodynamically more favourable, and therefore tighter, associations. For proteins capable of interacting with multiple partners within the same network context, ΔG values should be compared across all candidate interactions to identify the most energetically favourable interaction under a simplified competition framework. However, this needs to take into account several assumptions, including that the (i) comparisons are made under broadly comparable physicochemical conditions (i.e., pH, ionic strength, etc.); and most importantly, (ii) the effects of protein abundance, concentration-dependent occupancy, and kinetic constraints are to be neglected. Regardless, this feature of CAPRINI-M may provide a first-order approximation for prioritising dominant PPIs for downstream functional interpretation and experimental validation, as well as a ranking strategy for use in PPI-based network inference methods.

### 2.5 Neural Network (NN) - based PPI Predictions

Because leakage-minimised PPI benchmark datasets are currently most mature for human, we trained our NN-based predictors on a general (non-CVD-specific) human benchmark and then applied the resulting model to mouse PPIs in CAPRINI-M. To train NN-based machine learning models for PPI prediction, we used a previously published (Bernett et al. 2024), leakage-minimised general (non-CVD-specific) human PPI benchmark split into training/validation/test sets. For this, we used a balanced split of 2,439 protein pairs for training, 562 pairs for validation, and 608 pairs for testing (positives/negatives: Train 1,226/1,213; Val 279/283; Test 305/303) (Bernett et al. 2024). The selected high-confidence PPIs from DIGGER (Albrecht et al. 2025) were chosen based on their having complementary experimental information on the interaction/interfaces. Here, each protein pair was first modelled as a complex with AlphaFold3 and the top-ranked structure was retained based on AlphaFold3’s ranking score. From each predicted complex, a structured feature representation was built by combining direct AlphaFold3 confidence metrics (ipTM/pTM, minimum inter-chain PAE, fraction disordered, ranking_score) with estimated interface descriptors derived from the predicted geometry, including inter-chain contact statistics at multiple distance thresholds (3/5/7/10 Å) normalised by chain length and weighted by AlphaFold3’s contact-probability matrix, distance features from the closest inter-chain residue pairs (k=5/10/20), buried surface area fractions for each chain (Mitternacht 2016), and interface patch size/continuity features, that were complemented by established interaction-quality scores such as ipSAE (Dunbrack Jr. 2025), pDockQ (Bryant et al. 2022), pDockQ2 (Zhu et al. 2023) and LIS (Kim et al. 2024). For multimodal prediction, sequence and structure information were jointly encoded by concatenating embeddings from a pretrained protein-language model (ESM) (Villegas-Morcillo et al. 2022) with a selected set of AlphaFold3-derived features obtained from SHapley Additive exPlanations/SHAP analysis (top 12 AlphaFold3 features, SHAP12: ptm, bsa_a_norm_surface, chain_ptm_b, min_pae_a, pDockQ, ipsae_d0res_max, chain_ptm_a, biggest_joint_patch_10A, biggest_joint_patch_7A, biggest_patch_a_10A, biggest_patch_b_10A, bsa_b_norm_surface). A lightweight NN classifier was subsequently trained for interaction prediction, yielding probabilities. The model architecture, consisting of one or two fully connected hidden layers with ReLU activations and a single output neuron, was optimised. Learning rate, batch size, and epochs were jointly selected via 5-fold cross-validation, with the mean F1 score serving as the ranking metric. The final model was chosen through validation-based selection, and its performance was assessed on a held-out test set using accuracy, precision, recall, and F1 score.

Because both AlphaFold3-derived interface descriptors and ESM embeddings largely capture species-agnostic biophysical and evolutionary constraints, models trained on human interactions can generalise to mouse PPIs without re-learning mouse-specific sequence/structure priors. This is in line with recent cross-species evaluations, which report that PPI predictors trained on human data transfer successfully to mouse when built on protein-language-model representations (Liu et al. 2025; Li et al. 2025).

### 2.6 CAPRINI-M Web Application

CAPRINI-M (https://shiny.dieterichlab.org/app/caprinim) was developed using Shiny for Python and deployed with ShinyProxy (https://github.com/dieterich-lab/caprinim). The data layer consists of a PostgreSQL database and some non-tabular data on disk, which are queried via a read-only FastAPI endpoint from the Shiny frontend, enabling straightforward security controls and rate limiting. For the interactive molecular PPI visualisation, 3Dmol.js was used (Rego & Koes 2015). All other graphs were realised with Plotly. In the event of high traffic in the future, load balancing and replacing Shiny with a lighter alternative are possible improvements to the architecture.

### 2.7 Computational Benchmarking

To directly assess whether a cardiac-specific PPI resource provides added value over a generic interactome in systems-level analyses, we benchmarked pathway enrichment performance. We hypothesised that module reconstruction based on CAPRINI-M would generate disease/functional-associated modules whose interaction edges more accurately recapitulate established cardiac gene sets than modules derived from general PPI resources. We benchmarked pathway enrichment performance of three protein–protein interaction resources: CAPRINI-M, BioGRID (Oughtred et al. 2021), and physical protein interactions from STRING (Szklarczyk et al. 2022). For this, we relied on an edge–topology–based enrichment framework restricted to KEGG (Kanehisa & Goto, 2000), Reactome (Milacic et al. 2024) and Wikipathways (Kelder et al. 2012) in mouse. First, full interactomes for each resource were obtained, standardised to undirected gene–gene edges, and collapsed to unique canonical edge identifiers. Then, to mitigate bias, we retained those interactions in CAPRINI-M/BioGRID/STRING in which both interacting partners correspond to nodes present in the CAPRINI-M database. Next, KEGG/Reactome/Wikipathways pathway topologies were retrieved with the R package graphite-v2.10 (Sales et al. 2012). Pathways were then filtered to a cardio-relevant subset by keyword matching on KEGG/Reactome/Wikipathways pathway titles (i.e., contraction, cardiomyopathy, hypertrophy, ventricular, etc.). To define the benchmarkable reference set, pathways were required to contain at least 4 pathway nodes overlapping the benchmark gene universe and at least 3 pathway edges overlapping the union of edges across the three full interactomes.

For benchmarking, we relied on a recently published time-course multi-omics dataset from a murine model of progressive heart failure, TACOMA (Gjerga et al. 2024). The proteomics and gene expression data in the main TAC vs Sham comparison were jointly utilised to perform a network reconstruction analysis using the BioNet R-package (Beisser et al. 2010) to infer protein-protein interaction modules associated with heart failure progression. Edge-centric enrichment analyses were then performed against a resource-specific edge universe. Multiple testing correction across pathways was performed using the Benjamini–Hochberg (BH) procedure, and enrichment strength was summarised as −log10(BH-adjusted p value). Comparative benchmarking between CAPRINI-M and the two baseline resources (BioGRID and STRING) was then performed at the pathway level using paired t-tests and paired Wilcoxon signed-rank tests on matched pathway enrichment scores. All analyses and visualisations were implemented in R using tidyverse (data wrangling), ggplot2/patchwork (plotting), and graphite (pathway topology extraction).

### 2.8 Literature-Based Validation

To benchmark computationally predicted pairwise interaction favorability (using lower predicted ΔG as a proxy for stronger binding), we performed a targeted literature cross-validation for some of the reported PPIs against four experimental studies examining the relevant protein systems (Supplementary Table 1). For the HIF/ARNT system, we extracted reported PasB-domain binding thermodynamics and competition/affinity measurements comparing HIF-1α (HIF1A) and HIF-2α (EPAS1) interactions with ARNT from a biophysical/structural study employing ITC and AlphaScreen analyses (Cardoso et al. 2012). For the Notch ligand system, we evaluated direct binding measurements of the Notch1 ectodomain with DLL1 and DLL4, focusing on experimentally determined relative affinity differences between the two ligands (Andrawes et al. 2013). For the GJA1/Cx43 interaction network, we reviewed biochemical and cell-based competition data assessing c-Src (SRC) association with Cx43 and its effect on TJP1/ZO-1 binding (Toyofuku et al. 2001). Finally, for the BAG3 chaperone/co-chaperone system, we analysed in vivo and biochemical evidence for differential BAG3 dependence of HSPB8 versus HSP70/HSPA8-related sarcomeric/myofilament association, using these data as qualitative support for interaction selectivity in the absence of direct comparative thermodynamic measurements (Martin et al. 2021).

## 3. Results

To present the analyses supporting CAPRINI-M, we structured the Results to follow the main workflow steps. First, we benchmarked and selected prompting strategies for large language models (LLMs) to extract protein–protein interactions (PPIs) using the RegulaTome corpus. Next, we evaluated neural network–based predictors to assess how well AlphaFold3-derived features and protein language model embeddings distinguish true versus false PPIs. We then summarised the cardiac mouse PPI atlas, highlighting network properties and key annotations. Lastly, we tested the resource’s systems-level value by comparing pathway enrichment with general interactomes and validated whether predicted thermodynamic favorability matched experimental binding preferences in selected systems.

### 3.1 PPI RE Performance over RegulaTome

On the PPI RE task, LLaMA-3.3 70B’s performance varied substantially by prompting strategy and whether spaCy entity lists were provided (**Fig. 2**). The best-performing LLM-only configuration was *PosExamples*, reaching an F1 of 0.717 (95% CI 0.695–0.732) without pre-supplied entities. When spaCy-extracted entities were supplied to the LLM (LLM-spaCy), the best variant was AllExamples, with F1 = 0.686 (95% CI 0.669–0.703). Averaged across prompting variants, LLM-only runs showed higher precision but slightly lower recall (precision mean 0.769, recall mean 0.582) compared with LLM-spaCy (precision mean 0.716, recall mean 0.591), yielding similar mean F1 overall (LLM-only 0.660 vs LLM-spaCy 0.647). In other words, supplying entity lists tended to trade a modest decrease in precision for a small increase in recall, resulting in a negligible net change in F1 for PPIs. Across PPI prompting variants, precision was the most stable metric for LLaMA-3.3 70B (SD 0.028 in LLM-only; 0.018 with spaCy), whereas recall was the most sensitive to prompting (SD 0.083 in LLM-only; 0.049 with spaCy).

**Fig. 2.**
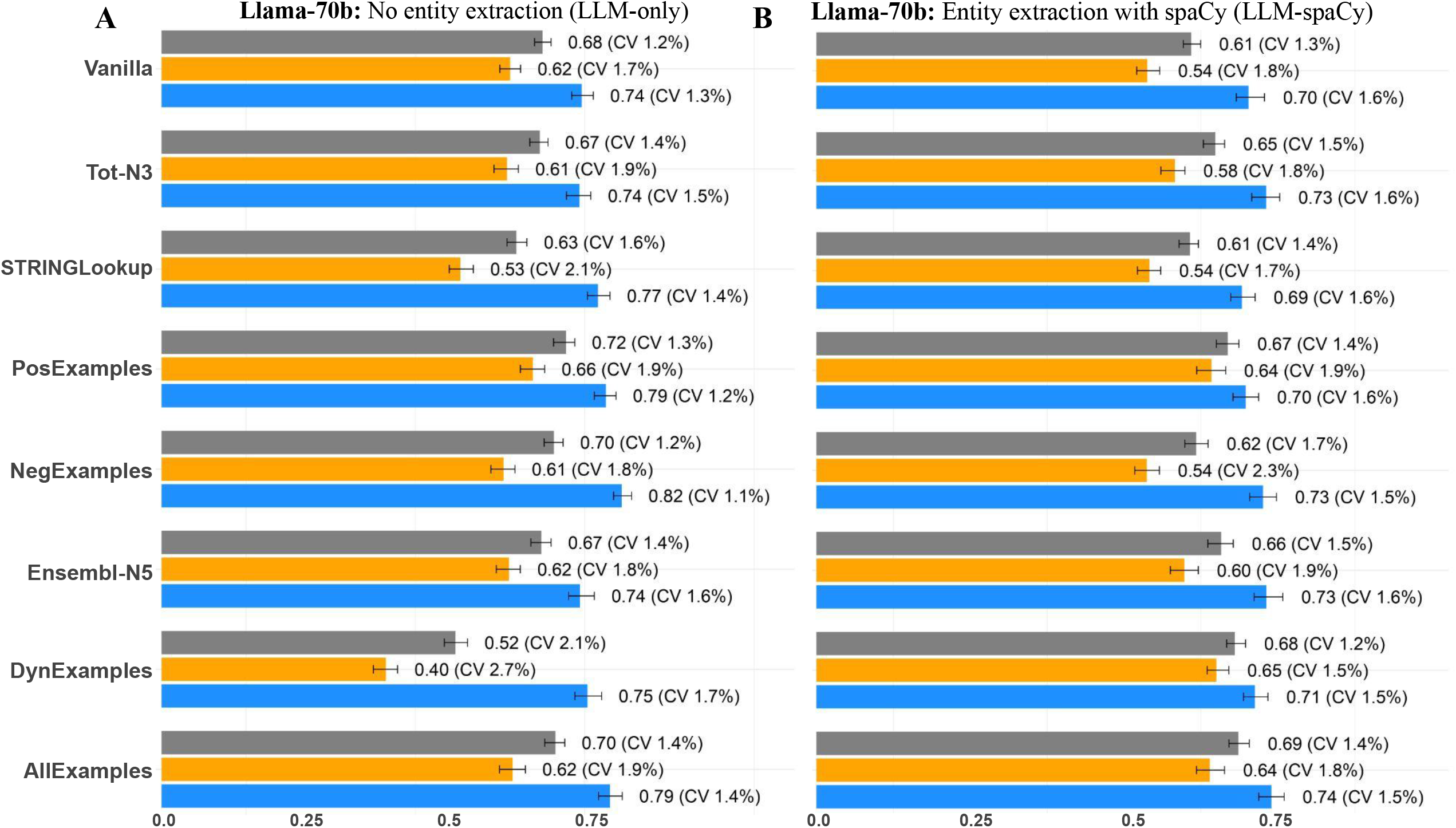
LLM-based evaluation of performance for the PPI-RE task (Precision – blue; Recall – yellow; and Grey – F1-score) across different prompting strategies when: **A**. No prior extracted entities were provided, and **B**. After providing entity names extracted with spaCy.

### 3.2 NN Performance

We evaluated all NN-based PPI predictors on the held-out test set of 608 protein pairs (305 positives, 303 negatives), ensuring that reported performance reflects generalisation to unseen proteins under leakage-minimised splitting (**Fig. 3**). Again, this evaluation was performed on a general (non-cardiac) human PPI benchmark dataset and is therefore independent of the cardiac corpus. Several models were tested, and details of each are reported in Supplementary Text 4. Across the tested approaches, the multimodal AF3+ESM models clearly performed best, with SHAP12_ESM3 achieving the top overall results in 4 out of 5 performance metrics (Accuracy = 0.890, Precision = 0.908, Recall = 0.869, F1 = 0.888, ROC-AUC = 0.961). Among sequence-only models, ESM3 was strongest (F1 = 0.868; ROC-AUC = 0.935) and showed the highest precision overall (0.910), while the best AF3-feature-only model (AF_all) reached solid but lower performance (F1 = 0.820; ROC-AUC = 0.908). Traditional baselines were competitive but behind the best deep/multimodal models (best SVM_SHAP12: F1 = 0.812; best RF_all: F1 = 0.763). Finally, baseline predictors such as D-Script underperformed mainly due to low recall (Recall = 0.50; F1 = 0.620), indicating that integrating AF3 interface/quality signals with ESM-3 embeddings yields the most accurate and well-balanced PPI predictions in this benchmark, as measured by ROC-AUC scores.

**Fig. 3.**
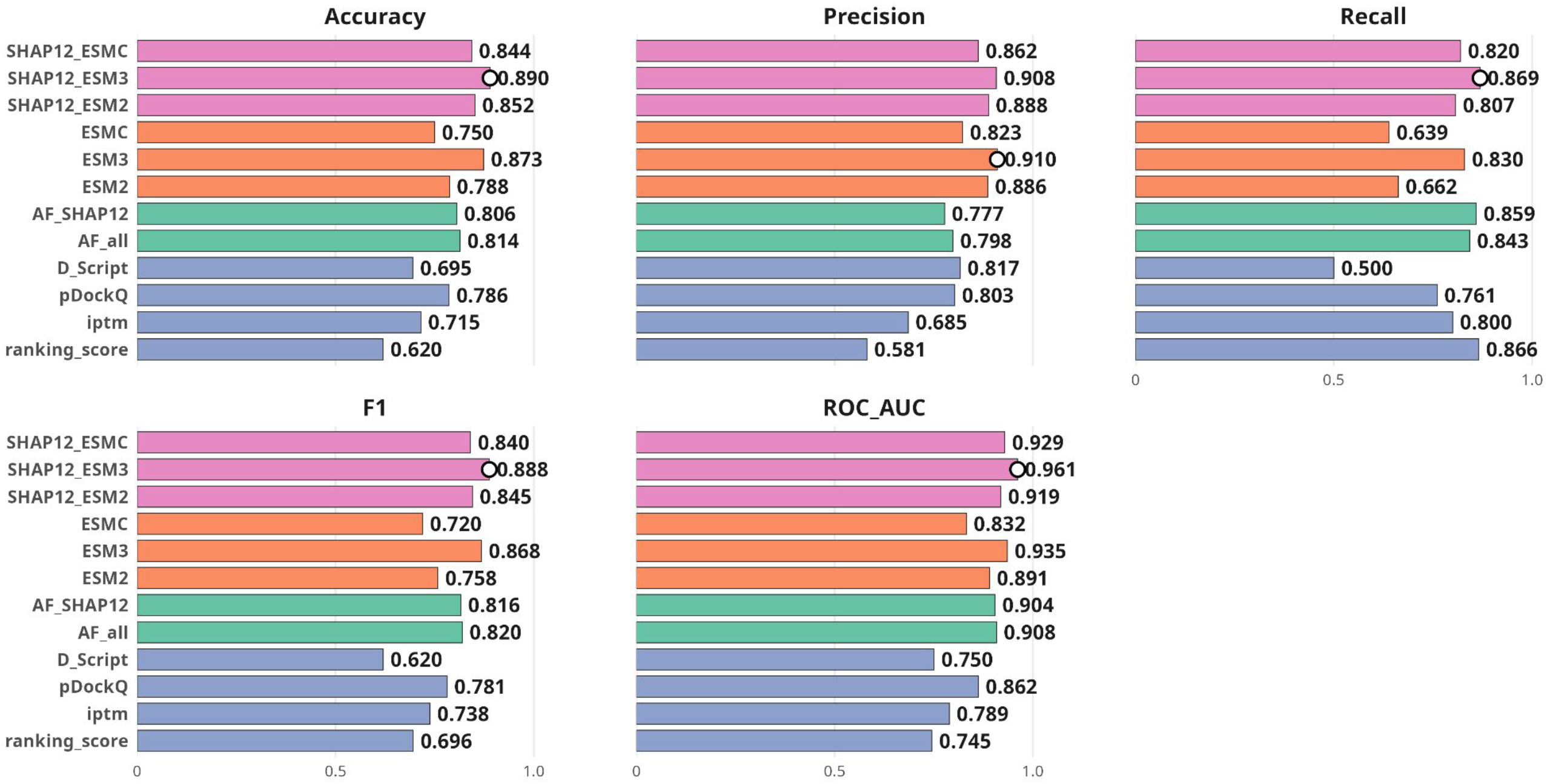
PPI prediction performance of multimodal AF3+ESM models (rosa bars); ESM-Sequence-only learned models (orange bars); Structural-only features (green bars); and baseline single-score confidences (blue bars). Information on the inputs used for prediction for each model is provided in Supplementary Text 4.

### 3.3 Cardiac PPI Atlas

The final Cardiac PPI corpus contains 11,189 interactions among 4255 unique proteins. Among these relations, 8,807 PPIs have only been reported in one study, while the remaining 2,382 PPIs have been reported in more than one study from the corpus. Among the 7,548 papers considered for extraction, only 4,503 reported at least one extracted PPI. Top-20 protein hubs in the network included: Nfkbiz, Tgfb1, Akt1, Stat3, Mapk14, Prkaa1, Itpkc, Pik3r1, Hif1a, Tnf, Nos3, Agtr1a, Trp53, Gsk3b, Sirt1, Mapkbp1, Nfatc3, Camk2a, Ctnnb1 and Tlr4. The average shortest path length between all pairs in the database is 4.168, while the longest shortest path between any protein pair is reported to be 12. When considering the best PPI classification model (AF3_SHAP12+ESM3), the average probability for the interactions to form a complex is 40.61% (SD: 25.69%), where 3,625 (32.40%) of PPIs report a complex formation probability higher than 50%. The average ΔG value reported across all PPIs is −20.74 kJ/mol (SD: 94.59 kJ/mol).

### 3.4 Benchmarking Analysis

Using the Reactome, KEGG and Wikipathways cardiac hypertrophy and contractile-focused edge-topology benchmark at the BioNet module edge FDR threshold of 0.05, 17 benchmarkable pathways (9 from KEGG, 6 from Reactome and 2 from Wikipathways) passed overlap/size filters (at least 4 nodes and at least three edges in the pathway set) and were evaluated across CAPRINI-M, STRING, and BioGRID solutions (**Fig.4-A**) (with 519, 3058 and 1729 BioNet module edges, respectively).

**Fig. 4.**
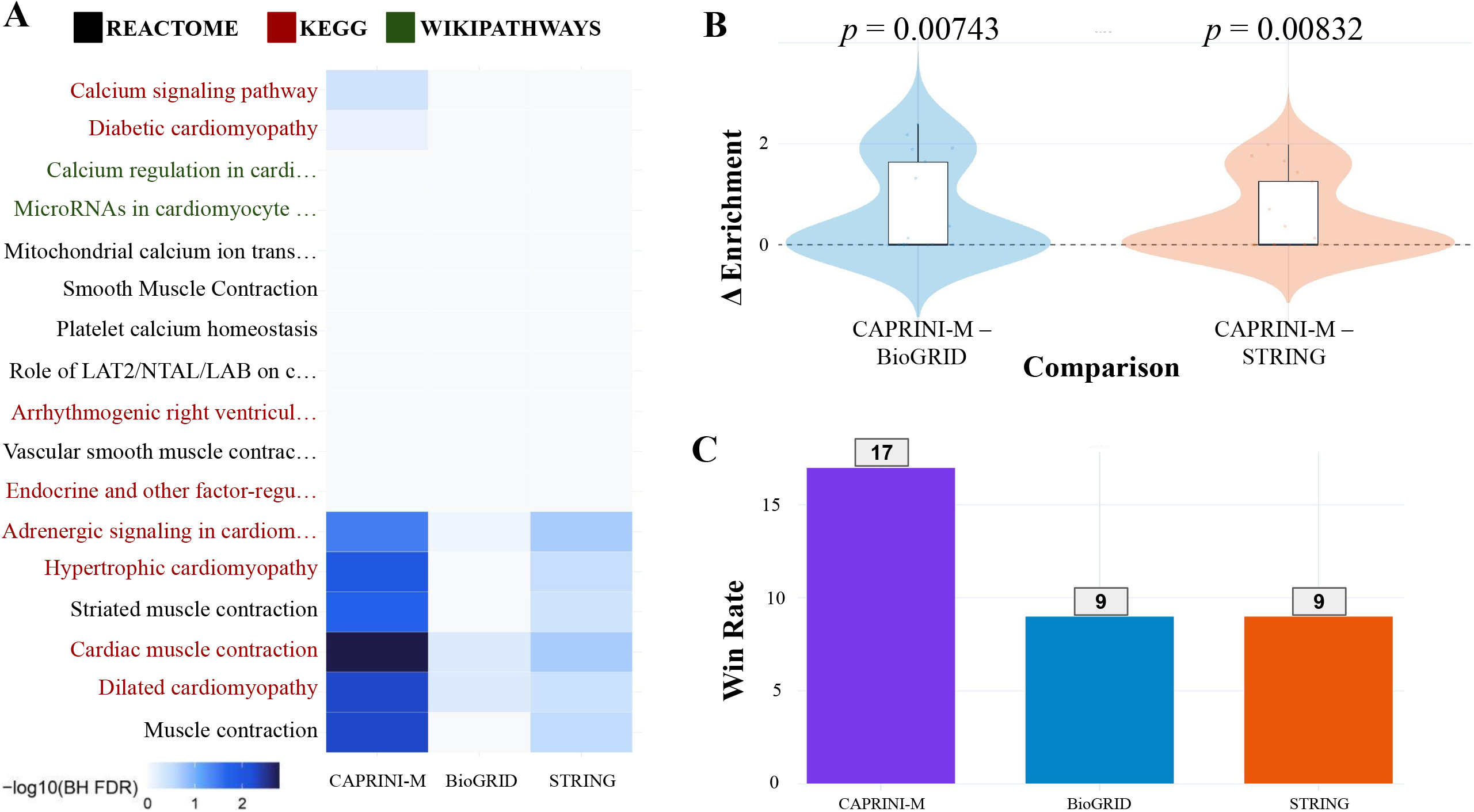
Benchmarking analysis results. **A**. Heatmap of −log10(BH-FDR) enrichment scores of cardio pathways across the BioNet network solutions analysed over the three PPI databases; **B**. Paired pathway-level differences (CAPRINI-M vs BioGRID and CAPRINI-M vs STRING) in enrichment scores combined with t-test statistics. **C**. Number of pathway wins by resource.

Across this combined pathway pool, CAPRINI-M showed the strongest overall enrichment (mean −log10(BH-FDR) = 0.73) compared with STRING (0.18) and BioGRID (0.04) (**Fig.4-B**). At BH-FDR < 0.1, CAPRINI-M identified 6 significant pathways, STRING identified 0, and BioGRID identified 0. Paired t-test pathway-level comparisons of −log10(BH-FDR) scores confirmed that CAPRINI-M enrichment scores were significantly higher than BioGRID (paired t-test: t = 3.06, p = 0.0074), while the difference versus STRING was weaker but still sensibly strong (t = 3.01, p = 0.0083) (**Fig. 4-B**).

Finally, win-rate analysis (most significant resource per pathway; ties counted for all winners) showed that CAPRINI-M achieved the most wins (17), followed by STRING (9) and BioGRID (9) (**Fig. 4-C**). The total exceeds 17 because 9/17 pathways had an FDR of 1. Among the 8 pathways with a unique winner, CAPRINI-M accounted for 8, with STRING and BioGRID each accounting for 0.

### 3.5 Experimental Support for Predicted PPI Rankings

Overall, the literature review supported our computational prioritisation framework in a tiered manner (Supplementary Table 1). This validation assesses not only whether the ΔG-based ranking aligns with experimental preferences, but also whether AlphaFold3 predicts interactions at biologically plausible domains or interfaces that correspond with the experimental systems under investigation (such as domain-restricted constructs, ectodomain binding assays, or competition and perturbation experiments). Experimental evidence was concordant with the computational ranking for the ARNT-centred HIF PasB interactions, as tighter binding of ARNT to HIF1A than to EPAS1/HIF2A was reported, consistent with a more favourable predicted ΔG[ARNT–HIF1A] = −410.39 kJ/mol relative to ΔG[ARNT–EPAS1] = −344.94 kJ/mol. In the GJA1/Cx43 system, activated SRC binding/phosphorylation was shown to reduce TJP1/ZO-1 association with Cx43 in cell-based and in vitro assays, qualitatively supporting a competitive preference consistent with ΔG[GJA1–SRC] = −3.58 kJ/mol being lower (more favourable) than ΔG[GJA1–TJP1] = 40.69 kJ/mol. Additionally, direct experimental affinity evidence supported NOTCH1–DLL4 (ΔG[NOTCH1–DLL4] = −217.23 kJ/mol) over NOTCH1–DLL1 (ΔG[NOTCH1–DLL1] = −68.26 kJ/mol), consistent with our lower predicted ΔG values for the more favourable pairs. Finally, for BAG3–HSPB8 (ΔG[BAG3–HSPB8] = −234.32 kJ/mol) versus BAG3–HSPA8/HSP70 (ΔG[BAG3–HSPA8] = −102.86 kJ/mol), the cited study (Martin et al. 2021) supports differential BAG3 dependence and subcellular targeting behavior (especially for HSPB8 at the myofilament/sarcomere) rather than a direct affinity ranking. For this particular case, our computational prediction suggests that the observed high apparent favorability of BAG3–HSPB8 may be due to thermodynamic factors beyond those initially highlighted (Martin et al. 2021). These could include more favourable interface packing/electrostatics, a smaller desolvation penalty, or a lower entropic cost upon complex formation, all of which would increase effective affinity under cellular chaperone assembly conditions. Therefore, while direct thermodynamic validation remains necessary, current experimental data support our predicted interaction hierarchy and warrant further quantitative biophysical testing.

A paired t-test was performed to compare predicted ΔG values for the favoured versus less favoured interactions across the four benchmarked cases, showing a significant difference in the expected direction (t = −3.86, p = 0.0307). These findings (although with a small sample size) support the capacity of our computational scoring framework to distinguish relatively more favourable PPIs, while underscoring the importance of expanding the validation cohort and incorporating direct quantitative binding measurements in future studies. Such findings establish CAPRINI-M as both a valuable resource for PPI prioritisation and a provider of mechanistically detailed interface and domain models. These models can inform targeted experimental validation and enhance the interpretation of competitive binding in cardiac signalling systems.

## 4. Conclusions & Discussions

In this paper, we have presented CAPRINI-M: an AI-curated cardiac-specific protein interactions that combines literature mining, structure-based analysis, and machine-learning prioritisation into a single accessible web resource. Starting from a large corpus of cardiovascular literature, we extracted a substantial set of cardiac-relevant PPIs and augmented these relations with AlphaFold3-derived structural/interface features, approximate thermodynamic descriptors, and interaction-likelihood scores. This design goes beyond conventional PPI repositories by coupling domain-specific curation with mechanistic annotations that can support hypothesis generation, interaction prioritisation, and downstream systems analyses in cardiovascular biology.

The CAPRINI-M framework demonstrated utility on multiple levels. First, prompt-engineered open-source LLMs enabled practical PPI extraction, enabling the scalable construction of a cardiobiology atlas. Second, multimodal PPI classifiers integrating AF3 and ESM features outperformed sequence/structure-only alternatives. Third, the resulting cardiac atlas showed improved cardio-focused pathway enrichment compared to general resources (STRING/BioGRID) and offered ΔG-based rankings concordant with experimental preferences. This ranking information is particularly useful for network inference analysis. Collectively, CAPRINI-M is a valuable resource for mechanistic cardiovascular systems biology analysis and translational target discovery.

Meanwhile, we also acknowledge some limitations of CAPRINI-M. For example, its coverage depends on accessible literature, meaning relevant findings in paywalled sources may be missed. Relation extraction, even with benchmarked prompting, is not flawless. AlphaFold3 modelling and feature extraction are computationally intensive. Additionally, ΔG estimates are approximations and should be interpreted with caution. Lastly, while ESM3 embeddings improve predictor performance, potential data leakage from pretraining on benchmark protein sequences means absolute performance should be interpreted conservatively. Dedicated benchmarking of interface accuracy remains a key future goal.

Future work will focus on enhancing both the context and coverage of CAPRINI-M. For example, we are currently developing a human version (CAPRINI-H) to broaden the resource’s applicability. On the text-mining front, we plan to expand the corpus of cardiobiology studies for deeper insights. Structurally, upcoming versions will integrate isoform-specific and PTM-aware modelling, explore alternative complex stoichiometries, and refine energetic and kinetic estimates to better capture the dynamics of cellular competition. In addition, we intend to conduct targeted benchmarking of predicted interaction interfaces against curated structural and biochemical datasets, and to apply CAPRINI-M’s interface-level data in splicing-aware network analyses. A key goal is to integrate CAPRINI-M with LINDA for future studies on single-cell cardiac datasets, leveraging LINDA’s strengths in combining expression data, differential splicing, and mechanistic constraints to map cell-type- and condition-specific network rewiring with greater resolution. Ultimately, CAPRINI-M will complement LINDA by providing comprehensive, literature-backed, interface- and stability-informed interaction hypotheses, with a workflow designed to be readily adapted for creating annotated interactome atlases in other tissues or disease domains beyond the cardiac field.

## Author Contributions

EG conceived the study, performed AF3 predictions and interaction and thermodynamic feature extraction, devised LLM aspects of the study, and supported the development of the PPI classification models. EG also supervised the work of CG and YZ. PW devised the LLM aspects of the study and worked on its implementation and evaluation. CG developed and deployed the web-application. YZ performed PPI feature extraction and devised and implemented PPI classification models. KP performed AF3 predictions used in the PPI classification models and contributed in its development. ML supervised KP and provided senior input on study direction and methodology. CD conceived and supervised the overall study and supervised EG, PW, CG, and YZ. All authors contributed to and approved the final manuscript.

## Acknowledgements

We would like to thank Harald Wilhelmi for the excellent infrastructure support.

## Availability

The CAPRINI-M web application is available online: https://shiny.dieterichlab.org/app/caprinim. The resource codes used for the shiny app development and deployment have been provided in GitHub: https://github.com/dieterich-lab/caprinim. Scripts used for the machine-learning models for protein–protein interaction prediction using AlphaFold3 complex outputs and derived structural features as well as computation of structural features have been provided on GitHub: https://github.com/AF3-PPI/AF3_PPI. The repository containing the information displayed in the CAPRINI-M web application, in tabular format, as well as the computational benchmarking analyses have been provided on GitHub: https://github.com/enio23/caprinim_ppi_info. Scripts for the LLM extraction-based evaluations and PPI RE extractions have been provided here: https://github.com/dieterich-lab/LLM_Relations. All the AF3 predictions have been provided in Zenodo: https://zenodo.org/records/18526268.

