## Supplementary Text for "CAPRINI-M: An AI-curated Cardiac-Specific Atlas of Protein Interactions in Mice"

Supplementary Materials:

Supplementary Text 1 2
Supplementary Text 2 2
Supplementary Text 3 5
Supplementary Text 3 5
Supplementary Table 1 6
Supplementary Figure 1 8

#### Supplementary Text 1 - RegulaTome Relation Annotation

RegulaTome interaction classes were categorised as PPIs as follows:

| PPI's |
|---|
| Complex Formation, Catalysis Of Posttranslational Modification, Catalysis Of Ubiquitination, Catalysis Of Methylation, Catalysis Of Deacetylation, Catalysis Of Phosphorylation, Catalysis Of Dephosphorylation, Catalysis Of Acetylation, Catalysis Of Glycosylation, Catalysis Of Acylation, Catalysis Of Deneddylation, Catalysis Of Small Protein Conjugation, Catalysis Of Sumoylation, Catalysis Of Other Small Molecule Conjugation Or Removal, Catalysis Of Demethylation, Other Catalysis Of Small Molecule Conjugation, Catalysis Of Adp-Ribosylation, Catalysis Of Palmitoylation, Catalysis Of Neddylation, Catalysis Of Phosphoryl Group Conjugation Or Removal, Catalysis Of Deubiquitination, Catalysis Of Small Protein Conjugation Or Removal, Other Catalysis Of Small Protein Conjugation, Other Catalysis Of Small Protein Removal, Catalysis Of Geranylgeranylation, Catalysis Of Farnesylation, Catalysis Of Lipidation, Catalysis Of Prenylation, Catalysis Of Small Protein Removal, Catalysis Of Deglycosylation, Catalysis Of Desumoylation, Catalysis Of Depalmitoylation, Catalysis Of Deacylation, Catalysis Of Small Molecule Removal, Other Catalysis Of Small Molecule Removal |

#### Supplementary Text 2 - Prompting Strategies

##### A. Base-prompt for PPI-RE

You are a top-tier molecular biologist specialised in the field of molecular biology. Following, you'll find a scientific TEXT, a desired OUTPUT FORMAT and a USER QUESTION. First, read the TEXT and study the OUTPUT FORMAT, then answer the USER QUESTION. {ner_list_prompt} Extract all the {interactions_type} interactions {ner_prompt} involved in signalling pathways from the text. Please only extract {target} pairs which directly interact with each other (i.e. through binding, phosphorylation, sumoylation, etc). Do not misinterpret functional relationships, co-occurrence, structural similarity, or indirect regulatory effects for direct interactions. {lookup_prompt}{dynex_prompt}.

**B. *PosExamples* for PPI-RE**

Below, you find some positive examples of protein-protein relations that give you an idea of what we are looking for:

P1: This cytokine induces p53 into a mutant-like conformation that forms a complex with Sp1
A1: p53 INTERACTS_WITH Sp1

P2: These findings suggest that the STAT3-NRF2 complex accelerates BLBC growth and progression by augmenting IL-23A expression.
A2: STAT3 INTERACTS_WITH NRF2

P3: HIF1A forms a transcriptional complex with ARNT under hypoxia.
A3: HIF1A INTERACTS_WITH ARNT

P4: PRMT1 methylates cGAS and suppresses cGAS/STING signalling in cancer cells
A4: PRMT1 INTERACTS_WITH cGAS

P5: TRAF6 ubiquitinates TGFβ type I receptor to promote its cleavage and nuclear translocation in cancer.
A5: TRAF6 INTERACTS_WITH TGFβ

P6: AKT1 phosphorylates AKT1S1 at Thr-246.
A6: AKT1 INTERACTS_WITH AKT1S1

P7: PIAS1 sumoylates PNKP in cells.
A7: PIAS1 INTERACTS_WITH PNKP

P8: CBP, but not p/CAF, acetylates GATA-1 at two highly conserved lysine-rich motifs present at the C-terminal tails of both zinc fingers.
A8: CBP INTERACTS_WITH GATA-1

P9: Experimental analysis revealed a chemical cross-linking between Hsp90 and the (CDK4)-cyclin D1 complex, stabilising their association during signal transduction.
A9: Hsp90 INTERACTS_WITH CDK4

P10: Similar to the CREB proteins, NFIX serves as a direct substrate of SRC1 and functions as a signal-responsive transcription factor.
A10: CREB INTERACTS_WITH SRC1;NFIX INTERACTS_WITH SRC1

P11: The N-terminal transactivation domain of p53, binds directly to the hydrophobic pocket of MDM2 to regulate its stability
A11: p53 INTERACTS_WITH MDM2

P12: Under physiological conditions, 14-3-3ζ exhibits weak, transient binding to Bad, allowing rapid modulation of apoptotic signalling in response to fluctuating

phosphorylation states.
A12: 14-3-3ζ INTERACTS_WITH Bad

P13: Inhibition of histone methyltransferase activity using the compound BIX-01294 diminished chondrogenic differentiation by downregulating cartilage-specific genes such as aggrecan and collagen type II.
A13: histone methyltransferase INTERACTS_WITH BIX-01294

#### C. *NegExamples* for PPI-RE

Below you find some examples of false positive protein-protein relations and the reason why you should not extract those:

P1: KRAS and BRAF cooperate in the MAPK signalling cascade to promote cell proliferation.
A1: Although the two proteins are in the same signalling system, the text does not provide evidence of a direct interaction.

P2: p53 and Protein MYC are both found in the same signalling complex.
A2: Incorrect assumptions based on co-occurrence or proximity.

P3: TNF and IL6 accumulate at DNA damage sites.
A3: Co-localisation, but no evidence of direct relation/interaction between the two.

P4: Gene TNF regulates the expression of Gene IL6.
A4: Misinterpretation of genetic or signalling pathways as protein interactions.

P5: Prmt5 shares 80% sequence identity with Protein Prmt7, which is known to bind BRAF.
A5: Incorrect assumptions based on structural similarity.

P6: PTEN was pulled down in a co-IP assay with CDKN2A.
A6: Incorrect interpretations of experimental methods.

#### D. Base-prompt for synonyms extraction

It has been noticed that in any of the LLM configurations, there was no guarantee that the entity names in the extracted relations would match exactly with how they were annotated in RegulaTome or spaCy. To address this issue, we additionally apply a prompt devised to extract 10 synonyms of each of the entities involved in the extracted relations. The prompt used for obtaining synonym names of the relation entities is: *“Generate at least 10 synonyms, generic names or abbreviations for the protein, gene or symbol provided by the user below.”*.

#### Supplementary Text 3 - Literature Filtering

Majority-voting three-prompt-based strategy for filtering non-cardiovascular studies from the fetched corpus:

1. You are an expert cardiologist and molecular cardio-biologist. Please verify whether this scientific paper is explicitly about heart function/disease, cardiovascular development, or vascular disease. Please only give me a Yes or No answer.

2. You are an expert cardiologist and molecular cardio-biologist. Please verify whether a major focus of this scientific paper is centred on cardiovascular/heart function and outcomes. Please only give me a Yes or No answer.

3. You are an expert cardiologist and molecular cardio-biologist. Please verify whether this scientific paper includes most of its content or at least a major section dedicated to the heart/cardiovascular domain. Please only give me a Yes or No answer.

#### Supplementary Text 4 - Tested PPI Prediction Models

##### **A.** Single-score AF3 confidence baselines

**pDockQ**: A single scalar derived from predicted interface size and confidence, commonly used to judge whether an AlphaFold(-Multimer/complex) interface is likely real. In our benchmark, it serves as a direct structural confidence baseline (higher = more plausible interaction).

**ipTM:** AlphaFold3's predicted interface TM-score, reflecting confidence in the relative placement/orientation of the interacting chains (higher = more confident interface).

**ranking_score:** AlphaFold3's internal ranking metric used to select the best of the five diffusion samples per job; here used both for "pick-top-model" and as a simple baseline score.

**D-SCRIPT:** An established sequence-based PPI predictor used as an external reference baseline (independent of your AF3 feature engineering pipeline).

##### **B.** Structure-feature learned models

**AF_all:** A neural-network classifier trained on the full engineered set of AlphaFold3-derived confidence + interface/geometry/contact features you extract from the top-ranked AF3 complex.

**AF_SHAP12:** The same type of NN classifier, but trained only on the top 12 AF3-derived features selected via SHAP feature-importance (a compact structure-only model).

**C.** Sequence-only learned models

**ESM2 / ESM3 / ESMC:** Neural-network classifiers trained on sequence embeddings from the corresponding pretrained ESM family model. No explicit AF3 structural features are used here (sequence-only predictors).

**D.** Sequence-only learned models

**SHAP12_ESM2 / SHAP12_ESM3 / SHAP12_ESMC:** Neural-network classifiers that concatenate ***(i)*** the SHAP-selected 12 AF3 structure/interface features with ***(ii)*** the respective ESM embedding, so the NN learns from both structural/interface evidence and sequence-derived representation.

#### Supplementary Table 1

| Target Protein (external gene name) | Proteins competing for the Target Protein (external gene names) | Winner protein more likely to bind (external gene name) | Study citation | Supporting sentence from the study |
|---|---|---|---|---|
| ARNT | HIF1A, EPAS1 | HIF1A | PMC3575918 | The HIF-1alpha/ARNT complex possesses a significantly higher binding affinity compared to the HIF-2alpha/ARNT complex |
| NOTCH1 | DLL4, JAG1 | DLL4 | PMC3757209 | Direct measurement of the binding affinity of the EGF6–15 region of Notch1 for Dll1 and Dll4 revealed that the Dll4 ectodomain binds with at least an order of magnitude higher affinity. |
| GJA1 | TJP1, SRC | SRC | PMID11035005 | Constitutively active c-Src |

| | | | | |
|---|---|---|---|---|
| | | | | inhibits endogenous interaction between connexin-43 and ZO-1 by binding to connexin-43 |
| BAG3 | HSPA8, HSPB8 | HSPB8 | PMC8134551 | Using western blot in myofilament-enriched LV tissue from wild-type, BAG3+/−, and BAG3−/− mice, we found myofilament levels of HSPB8 and CHIP were reduced in the partial and complete absence of BAG3 (Fig. 5a–d). However, myofilament HSP70 expression was not impacted by decreased BAG3, suggesting it requires different mechanism for targeting to the sarcomere. |

### Supplementary Figure 1

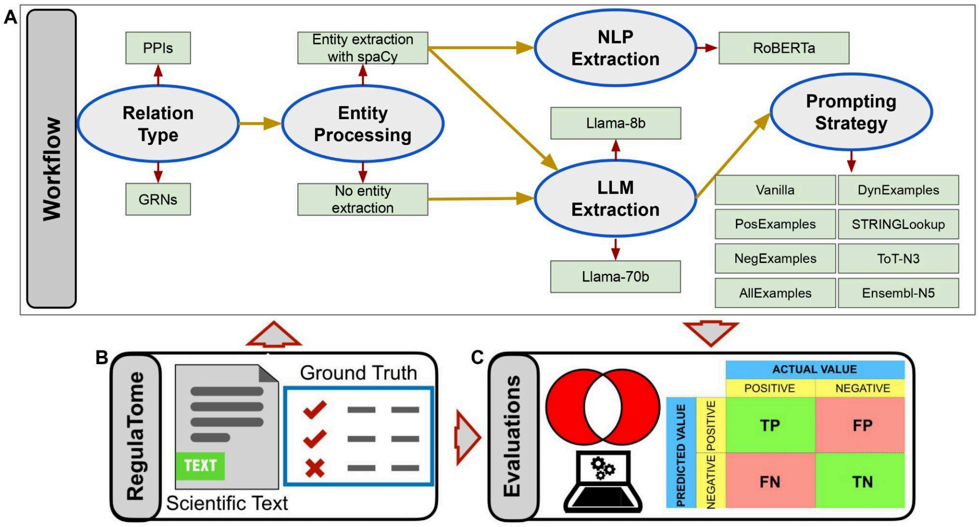
